# Synapses without tension fail to fire in an in vitro network of hippocampal neurons.

**DOI:** 10.1101/2023.06.24.546401

**Authors:** Md Saddam Hossain Joy, Duncan L. Nall, Basher Emon, Ki Yun Lee, Alexandra Barishman, Movviz Ahmed, Saeedur Rahman, Paul R. Selvin, M. Taher A. Saif

## Abstract

Neurons in the brain communicate with each other at their synapses. It has long been understood that this communication occurs through biochemical processes. Here, we reveal a previously unrecognized paradigm wherein mechanical tension in neurons is essential for communication. Using *in vitro* rat hippocampal neurons, we find that (1) neurons become tout/tensed after forming synapses resulting in a contractile neural network, and (2) without this contractility, neurons fail to fire. To measure time evolution of network contractility in 3D (*not* 2D) extracellular matrix, we developed an ultra-sensitive force sensor with 1 nN resolution. We employed Multi-Electrode Array (MEA) and iGluSnFR, a glutamate sensor, to quantify neuronal firing at the network and at the single synapse scale, respectively. When neuron contractility is relaxed, both techniques show significantly reduced firing. Firing resumes when contractility is restored. Neural contractility may play a crucial role in memory, learning, cognition, and various neuropathologies.

## Introduction

A critical function of neural networks, involving learning, memory, and computation, is to link to the firing pattern of the neurons involved ^1^. Loss of firing has been associated with several neurodegenerative diseases such as Alzheimer’s disease, Dementia, Huntington’s disease, Amyotrophic Lateral Sclerosis (ALS), and Parkinson’s disease, among others ^2^. In Alzheimer’s disease, for example, there is evidence that neuronal activity decreases long before the degeneration of neurons and loss of synapses ^3^. Thus, determining the various factors that regulate neuron firing is important in understanding neurodegenerative diseases and related processes.

Neuron firing has long been understood as regulated by biological and chemical processes (^4–6)^. Mechanical force or tension, commonly associated with muscle, has never been considered as a part of neural function or firing. There is, however, growing evidence suggesting that neurons are mechanosensitive. For instance, externally applied tension or pull induces axonal growth ^7, 8^; mechanically stretched brain slices exhibit hyperexcitability ^9^; applied stretch on Aplysia axons modulates vesicle transport ^10^; stretch on frog leg neurons increases neurotransmitter release ^11^. In addition, neuronal tension is required for the synaptic vesicle clustering at neuromuscular junctions in Drosophila fruit fly embryos, which is a prerequisite for successful neuron firing ^12^. These findings led us to hypothesize that mechanical tension in mammalian brain neurons may also influence neuron firing, in addition to other biochemical factors. We have chosen to examine this possibility using primary rat (mammalian) hippocampal cells due to their role in learning and memory ^13^, as well as in neurodegenerative diseases ^14–16^. Understanding the role of mechanical tension in such neurons may lead to novel therapies for neurodegenerative diseases exploiting tension as a target.

We cultured primary rat hippocampal neurons in a 3D extracellular matrix (ECM). Neurites from these neurons extend into the ECM and form synapses or junctions with each other. This results in an interconnected network of neurons. Neurites tend to shrink after synaptogenesis, i.e., they become taut or mechanically tensed. This makes the network to become contractile when neurons adhere to the ECM or a substrate where they remain anchored. We measured the emerging force of the network of hippocampal neurons as they form synapses with time. Over the past few decades, various methods for measuring neuronal force have been developed, including Vectorial Analysis ^17^, Traction Force Microscopy (TFM) ^18^, Optical Tweezers ^19^, Microneedles ^20, 21^, and Atomic Force Microscopy (AFM) ^22^. However, none of these methods can measure neuron forces in a 3D matrix; neither can they quantify forces as neurons form synapses. To our knowledge, there is currently no technique available that can measure the force produced by a network of neurons in 3D. To address this gap, we developed an ultra-sensitive force sensor, made from PDMS by molding, with 1 nN resolution, capable of measuring the force of a neuronal network ^23^ and applied it to rat hippocampal neurons. We asked: what is the time evolution of network force as the neurons form synapses; whether the neurons maintain mechanical tension after synaptogenesis; whether neuron force can be modulated (relaxed) with drugs; whether neurons regain contractility after drug washout; and the effect of force disruption on neuronal firing rate and presynaptic vesicle clustering.

We used Multi-Electrode Array (MEA) and iGluSnFR ^24^, a glutamate sensor, to quantify neuronal firing at the network scale and at the single synapse scale, respectively. Our results demonstrate that the hippocampal neurons indeed maintain mechanical tension after synaptogenesis. Disrupting this tension using Blebbistatin (which inhibits Myosin II-induced contractility) reduces both the firing rate and amount of presynaptic vesicle clustering of hippocampal neurons. Restoring tension with Blebbistatin-washout recovers firing. The development of the force sensor capable of measuring the force produced by the neuronal network in 3D may be a valuable tool for future studies in neuroscience.

## Materials and Methods

### Rat hippocampal neuron extraction and culturing

Primary rat hippocampal neurons were harvested from day 18 embryonic (E18) rat hippocampi using the standard protocol ^25^. Briefly, hippocampi from the E-18 embryo brain were dissected in slice dissection solution (SLDS) containing 82 mM Na_2_SO_4_, 30 mM K_2_SO_4_, 10 mM HEPES, 10 mM Glucose, and 5 mM MgCl_2_. Dissected hippocampi were then digested in 3mg/ml protease 23 (Sigma P4032) in SLDS for 10 minutes. After that, a fire-polished glass pipet with a small opening was used to dissociate hippocampi into single cells. Dissociated rat hippocampal cells were plated on 25 mm poly-l-lysine-coated glass coverslips and cultured at 37 °C with 5% CO_2_ in neurobasal media (ThermoFisher, 21103049) supplemented with B27 (GIBCO), 2uM Glutamax, 50 unit/ml penicillin, and 50 unit/ml streptomycin. Half of the medium was replaced with fresh maintenance medium twice a week. At 13/14 days in vitro (DIV), cells were co-transfected with Homer-miRFP/Lifeact-miRFP (1ug/coverslip) and iGlusnFR (1ug/coverslip) using lipofectamine 2000 transfection reagent. After 48 hours, coverslips were transferred to an imaging chamber (Warner RC-40LP) containing HEPES-buffered saline (140 mM NaCl, 5 mM KCl, 25 mM HEPES, 10 mM D-glucose, 1.5 mM MgCl_2_, and 2 mM CaCl_2_ at pH 7.4) for imaging.

### Fabrication of PDMS sensors

The detailed method of fabrication of the PDMS sensors is described elsewhere ^23, 26^. Briefly, a polydimethylsiloxane (PDMS) mixture was prepared by mixing polymeric base and curing agent (Sylgard 184) at a 10:1 ratio by weight. This liquid PDMS mixture was then poured into a microfabricated silicon wafer mold with a nominal depth of 200 um to cast the sensors. Capillary tension helped the liquid PDMS to reach all the micro features and trenches. The filled-up silicon wafer mold was placed in a 60°C oven overnight (∼12 hours) to cure the PDMS. The sensors were then carefully lifted off the silicone mold using fine tweezers and afterward autoclaved to sterilize them. An SEM image of the sensor is shown in Figure 1(a) and a simplified view of the neuronal force measurement is shown in Figure 1(b).

**Figure 1.**
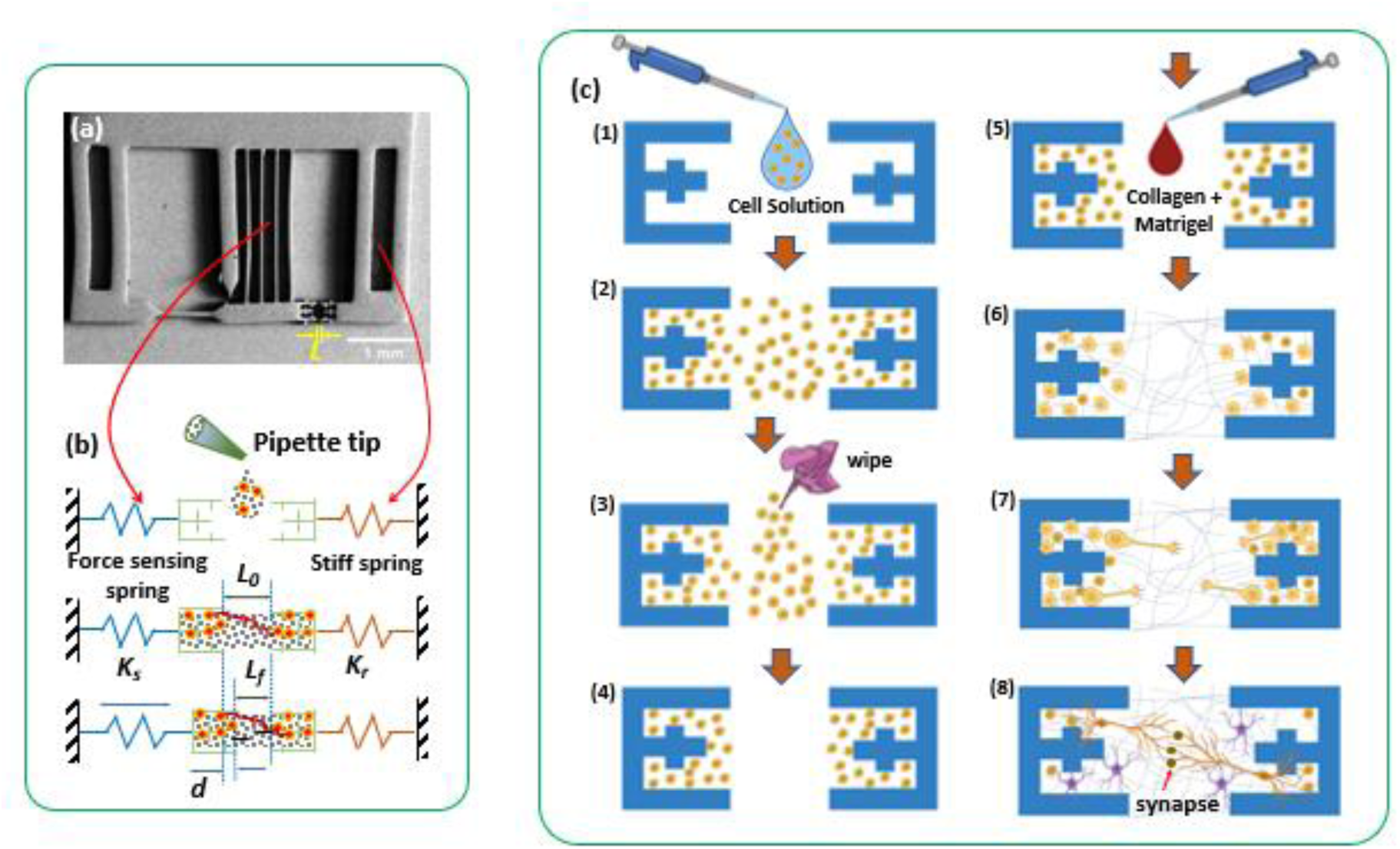
Schematic representation of the concept and methodology to measure neuronal force using the ultrasensitive force sensor. (a) SEM image of the force sensor; (b) the simplified view of neuronal force measurement technique in a 3D matrix (adapted from Emon et al. 2021); (c) The schematic diagram of the steps to form the 3D neuronal tissue

### Experimental setup and ECM solution

The experimental setup of the force sensors was done following the protocol as described by ^26^ with slight modifications. Briefly, a gelatin layer was formed in the well of the glass bottom petri dish. Then the force sensors were placed on top of the gelatin layer with its base attached to a double-sided tape. All the space between the beams was filled with gelatin. The gelatin layer works as a sacrificial layer. It gives support to the sensor during the tissue formation and helps to avoid the formation of a bubble that can affect the force measurement of the sensor. Gelatin was melted at 37°C and washed out after the tissue formation.

The extracellular matrix (ECM) solution mixture containing the Collagen and Matrigel was freshly prepared on ice. At first, type-I collagen from rat tail (Corning) was neutralized with 1N sodium hydroxide, 10X PBS, and deionized (DI) water. Then Matrigel (Corning) was mixed with neutralized collagen to get a final concentration of 1mg/ml of each. This tissue precursor solution was kept on ice and put in a 4°C chamber of the fridge till used for tissue formation.

Figure 1(c) illustrates the detailed method of 3D neuronal tissue formation between the two grips of the force sensor. To maintain a cell-free area in the middle region between the grips, at first, we pipetted high-density cell suspension (∼10 million/ml) of rat hippocampal neurons on top of the grips. Usually, capillary tension was enough to draw the cell suspension inside the grip and drive the air out. However, low-pressure vacuuming for about 30-40 seconds using a vacuum desiccator was occasionally used to remove persistent bubbles. Wiping was used to carefully suck out any cell suspension from the space between the two grips. The ECM solution without any cells was pipetted on top and between the grips forming a capillary bridge and kept at room temperature for about 40-50 minutes to polymerize and form the tissue. The resulting tissue consisted of two ends with neurons and ECM anchored by the sensor grips, while space in between them with only ECM. We expected the neurites to approach this space and form synapses. Any contractile force of the neurites is sensed by the sensor.

### Glial treatment

In order to measure the force of neuronal tissue in the absence of glial cells, a glial cell suppression protocol was performed based on the method described by ^27^. On day 1, the sample was treated with a cocktail of 20 μM Fuorodeoxyuridine (FUDR) (MP Biomedicals), 20 μM uridine, and 0.5 μM Ara-C (all from Sigma) for 72 hours to suppress the glial cells from the tissue. After 72 hours, two-thirds of the cocktail solution was replaced with neuron culture maintenance media. The cultures were then maintained by re-feeding every 3 days with neuron culture maintenance media, with half of the medium being replaced with fresh maintenance medium.

### Immunohistochemistry and confocal imaging

For immunofluorescence imaging, the samples were fixed with 4% Paraformaldehyde (PFA) in Phosphate buffered saline (PBS) for 1 hour. Subsequently, 0.2% Triton X-100 in PBS was used to permeabilize the samples, and 2.5% Bovine Serum Albumin (BSA) with 2% normal goat serum (NGS) in PBS was used as a blocking solution. Samples were then incubated overnight in primary antibody solution at 4°C. The next day, samples were washed with PBS five times, 5 minutes each, and then incubated in secondary antibody solution for 2 hours at room temperature. Afterward, the samples were washed with PBS three times, then incubated in 4′,6-diamidino-2-phenylindole (DAPI) (1:1000) (Invitrogen, Carlsbad, CA, USA) for 10 minutes and washed with PBS again. A confocal microscope, LSM 710, with an EC Plan-Neofluar 20X/0.5 N.A. objective lens (Carl Zeiss AG, Oberkochen, Germany) was used for the image acquisition. Maximum intensity projection of the acquired confocal z-stacks was constructed using the ImageJ (U.S. National Institutes of Health, Bethesda, Maryland, USA) software, and IMARIS (version 9.6.0, Bitplane AG, Zurich, Switzerland) was used for the spot reconstruction.

### Imaging and Image analysis

An inverted optical microscope (Olympus IX81, 20X objective, Olympus America Inc., Center Valley, PA) was used for Bright field time-lapse imaging. The microscope was mounted on a vibration isolation table (Newport Corporation, Irvine, CA) and it had an environment-controlled chamber. The chamber maintained cell culture conditions at 37 °C temperature, 5% CO2, and 70% humidity. A Neo sCMOS camera (active pixels 1392 × 1040, with a pixel size of 167 nm) (Andor Technology, Belfast, Northern Ireland) was used to acquire the image of both the tissues and the gauges. Displacements of the sensor gauges were measured using the template matching plugin in ImageJ with sub-pixel resolution ^23^.

Fluorescence images of neurons firing (using iGluSnFR) were acquired with a Nikon Eclipse Ti microscope with a Nikon Apo TIRF 100× objective (NA 1.49). The sample was stabilized in the z-axis using the Perfect Focus System (PFS) on the microscope and illuminated with an Agilent laser system (MLC400B) with four fiber-coupled lasers (405, 488, 561, 640 nm). Images were recorded using a back-illuminated EMCCD camera (Andor DU897). A motorized stage with a piezo-top plate (ASI PZ-2000FT) was used for x−y−z control. Elements software (Nikon) was used for image acquisition.

Detection of spikes from the fluorescence signal of iGluSnFR was performed using a modified version of the method described by ^28^. Specifically, filtered raw fluorescence traces were extracted from each individual synapse and analyzed for peak detection. Successful glutamate release events were identified as peaks with amplitudes two standard deviations above the average of the baseline trace.

### MEA preparation, recording, and spike/burst detection

MEA recordings were performed using a MEA 2,100-Lite Amplifier (Multi Channel Systems MCS GmbH) and a 6-well MEA device fabricated by the same manufacturer. The device contained 9 embedded 30 μm diameter TiN electrodes per well with 200 μm spacing between electrodes and six reference electrodes. Prior to cell culture preparation, the MEA platform was coated with 0.1 mg/ml Poly-D-Lysine (Sigma) and Matrigel. MEA measurements were carried out at 37 °C for 5 minutes at a sampling rate of 10 kHz with a sealed cover to maintain CO2 concentration. Media was replenished every other day. Neuronal activity was analyzed by Multi-Channel Analyzer software (Multi Channel Systems MCS GmbH), Python (3.9.7), and MATLAB. Raw data were filtered using a 2nd order Butterworth high pass filter with a 200 Hz cutoff frequency. Action potentials were detected as spikes by a threshold of 4 × standard deviations for both rising and falling edges from the noise magnitude distribution. Only active electrodes containing at least 5 spikes/min were considered for spike detection. Bursts were defined based on criteria such as a minimum of 4 spikes, lasting at least 50 ms, separated from another burst by at least 100 ms, and the interval between the first two and the last two spikes of a burst should be less than 50 ms, respectively.

### STORM imaging

Hippocampal neurons were treated with blebbistatin or vehicle for 30 minutes at 16 DIV with 4% PFA/sucrose. We permeabilized with .5% Triton X supplemented with 5% BSA for 30 minutes. We stained the primary antibody against synaptophysin (Abcam) (1:500) overnight at 4°C. We washed 3x with PBS and stained the primary with an alexa647 labeled secondary at room temperature for 1 hour. Samples were washed 3x with PBS and taken for imaging.

STORM imaging was done on a Nikon TI Eclipse equipped with a 1.49 NA 100x oil immersion objective. All imaging was done using NIS-elements software. Lasers (405, 488, 561, 640nm) were passed through an AOTF and fed into a single-mode optical fiber housed in an Agilent laser system (MLC400B). The lasers were collimated and focused on the back aperture of the objective via a TIRF module. The lasers were bounced off a quad-band dichroic (Chroma, ZT405-488-561-640RPC) and imaged in a HILO configuration. Axial stability was maintained with Nikon’s perfect focus system. Collected fluorescence was directed through an additional 1.5x magnifying lens. The fluorescence passed through a filter wheel equipped with various bandpass filters (525/50, 600/50, 680/40) and was imaged onto an EMCCD (DU897, ANDOR). We used the AOTF to pulse 405 for 100ms followed by the acquisition of 4 frames of fluorescence imaging (100ms exposure). To correct for lateral focus, fiducial beads were imaged by an IR LED (760nm) onto a secondary camera (DMK 23U274, The Imaging Source). One frame of lateral drift correction was imaged for every four frames of fluorescence. STORM imaging was done in the presence of an imaging solution containing 10% oxyrase (Oxyrase, Inc), 100 mM cysteamine, and 20 mM sodium DL-lactate in PBS.

## Results

### Network of neurons maintain mechanical tension after synapse formation

#### Principle of force sensor

The fabrication and principle of operation of the force sensor for measuring the force of single or multiple cells were discussed in ^26^. Here, we modify the cell plating process on the sensor to measure neural network force. The sensor comprises of a soft spring, a stiff spring, and two grips, attached to the springs, that secure a self-assembled tissue. The SEM image of the sensor and its working principle are depicted in Figures 1(a) and 1(b), respectively. The soft spring, which serves as the force-sensing component, is represented by four thin mechanical beams, while the stiff spring is represented by thick beams. The stiffness of the soft and rigid beams is represented by K_s_ and K_r_, respectively. The neuronal tissue is situated between the grips, with the cell body of neurons positioned within the grips (Figure 1(c)). As the neurons extend their neurites towards the opposite grip through the ECM bridge, the sensor records the force of the growth cones collectively. Upon the formation of synapses, the neurites contract and generate contractile (pulling) forces, which the sensor captures as a time-dependent measurement of post-synaptogenesis force. The displacement, d, of the force-sensing beams at any given time point, t, is calculated as d = L_0_ - L_f_(t), where L_0_ is the initial gap between the grips and L_f_(t) is the gap at that time *t*. Consequently, the force produced by the neuronal tissue at *t* is given by F(t) = K_s_ * d(t), where the time after ECM curing is considered as t=0. Displacements, d(t), are measured from time-lapse optical images of the sensor.

#### Neurons grow and form mature synapses on force sensor

We first validated that rat hippocampal neurons grow and form mature synapses in the tissue formed on our force sensor (Figure 2, Supp videos 1 and 2). Figure 2(a) presents bright-field images of the space between the two grips at various time points. Soon after gelling of the ECM bridge, the space between the grips was devoid of any cell bodies or neurites. However, at DIV2 of the culture, neurites became visible in the middle space (indicated by the arrows). With time, more neurites became visible. To confirm the presence of mature synapses, the sample was fixed at DIV14 and stained for the neurite marker MAP2 (cyan), presynaptic marker Bassoon (green), and post-synaptic marker Homer (red) (Figure 2(b)). Immunofluorescence images show that the pre-and post-synaptic markers were along the neurite and colocalized with each other, indicating the presence of mature synapses. The co-localization of pre-and post-synaptic markers was further confirmed by reconstructing the presynaptic Bassoon and postsynaptic Homer signal using the spot feature of IMARIS (Figure 2(c)). The IMARIS-reconstructed image clearly showed that the presynaptic marker (Bassoon, red) and post-synaptic marker (Homer, green) were adjacent to each other, confirming that the rat hippocampal neurons were forming mature synapses in the 3D tissue on the sensor platform.

**Figure 2.**
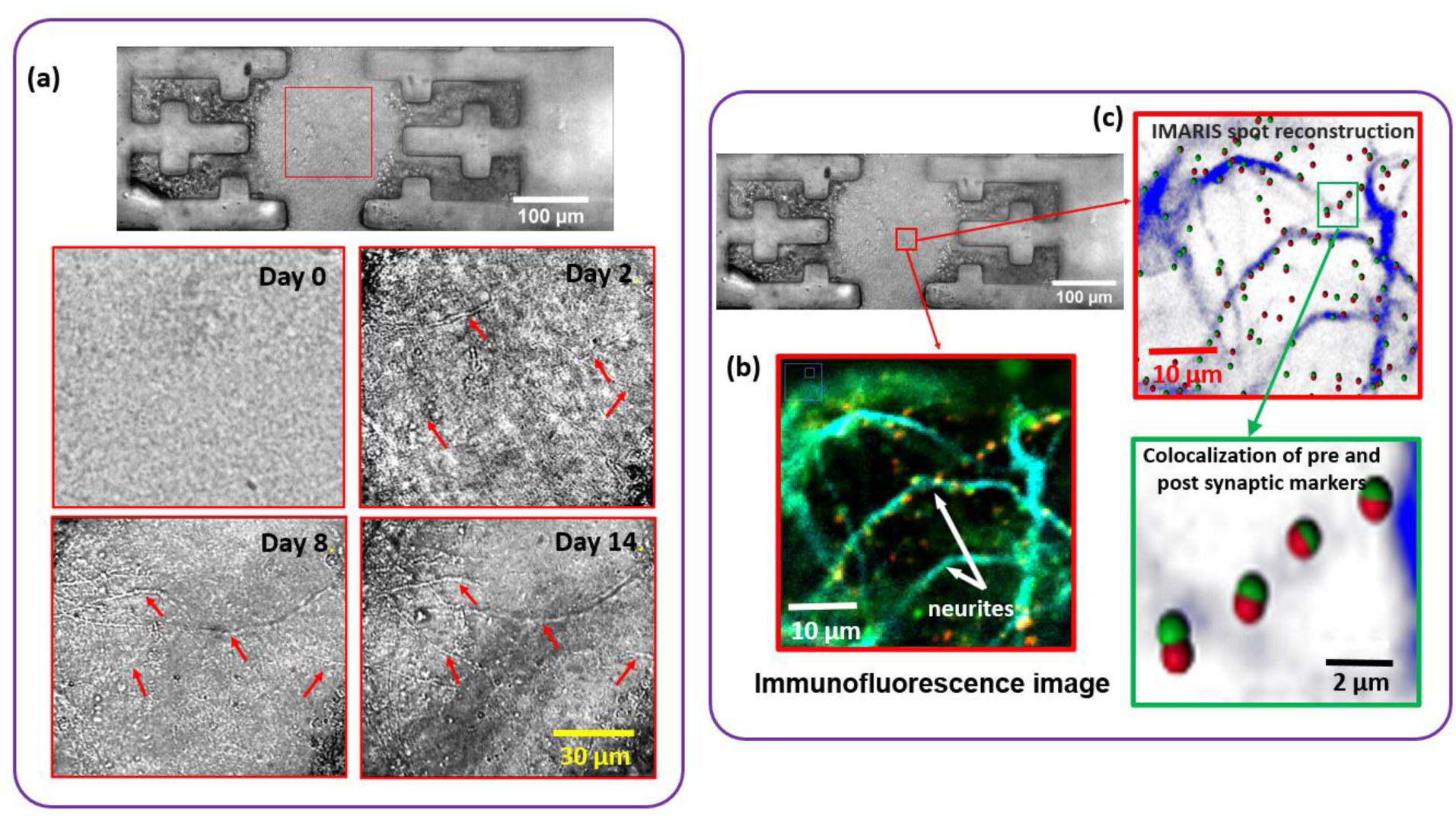
Rat hippocampal neuron grow and form mature synapse in the force sensor platform. (a) Bright-field (BF) images of neuronal tissue. The zoomed-in view shows the growth of neurite (indicated by arrow) at different time points; (b) Immunofluorescence image of neuronal tissue (neurite marker MAP2 (cyan), Presynaptic marker Bassoon (green), and post-synaptic marker Homer (red).) and (c) the corresponding IMARIS spot reconstruction confirms the colocalization of pre and posts synaptic markers and formation of the mature synapse.

#### Measurement of force produced by hippocampal neuron in 3D matrix

The temporal variation of the neuronal network force of several samples was measured using the force sensor (Figure 3(a)). Measurement started 24 hours after neuron plating. By that time, the sacrificial gelatin layer (see Methods) was completely dissolved in the media. The Force of the neuronal tissue was measured every 5 minutes, for 24 hrs., (Figure 3(b)). Based on the tracking of the growth cones, the average velocity of neurite advancement (measured since the plating of neurons) was found to be in the range of 0.5 to 0.9 μm/min (Figure S6). The gap between the two grips of the force sensor was 100 μm, implying that within 24 hours of plating and before force measurement started, most of the neurites have traveled the gap between the grips and might have contacted each other forming synapses. Therefore, the force recorded was mostly during synaptogenesis. The bright-field image in Figure 3(a) depicts the displacement of the grip attached to the soft spring. Displacement increases due to the contractile force produced by the neurons. For the three samples shown (Figure 3(b)), neurons gradually increase their force. The rate of force increase was approximately 0.5 nN/hr. Notably, the force generation pattern of neurons differs significantly from that of fibroblasts ^23^. Fibroblasts exhibit fluctuations in force over time, with peaks and drops in the force profile resulting from the cells’ continuous contraction and relaxation ^23^. Neurons display a continuous increase in force with time with minimal fluctuation. Neurons in vitro need about 14 days to form functional synapses ^29^. Hence, we measured network force for a total of 14 days, every 5 mins the first day and once a day the rest 13 days (Figure 3(c)). Recall that force measurement started after 24 hrs. of cell plating. For most of the samples, neurons continued to increase the force for up to 6 to 8 days and then reached saturation. The maximum force varied between samples within 20 nN to 65 nN.

**Figure 3:**
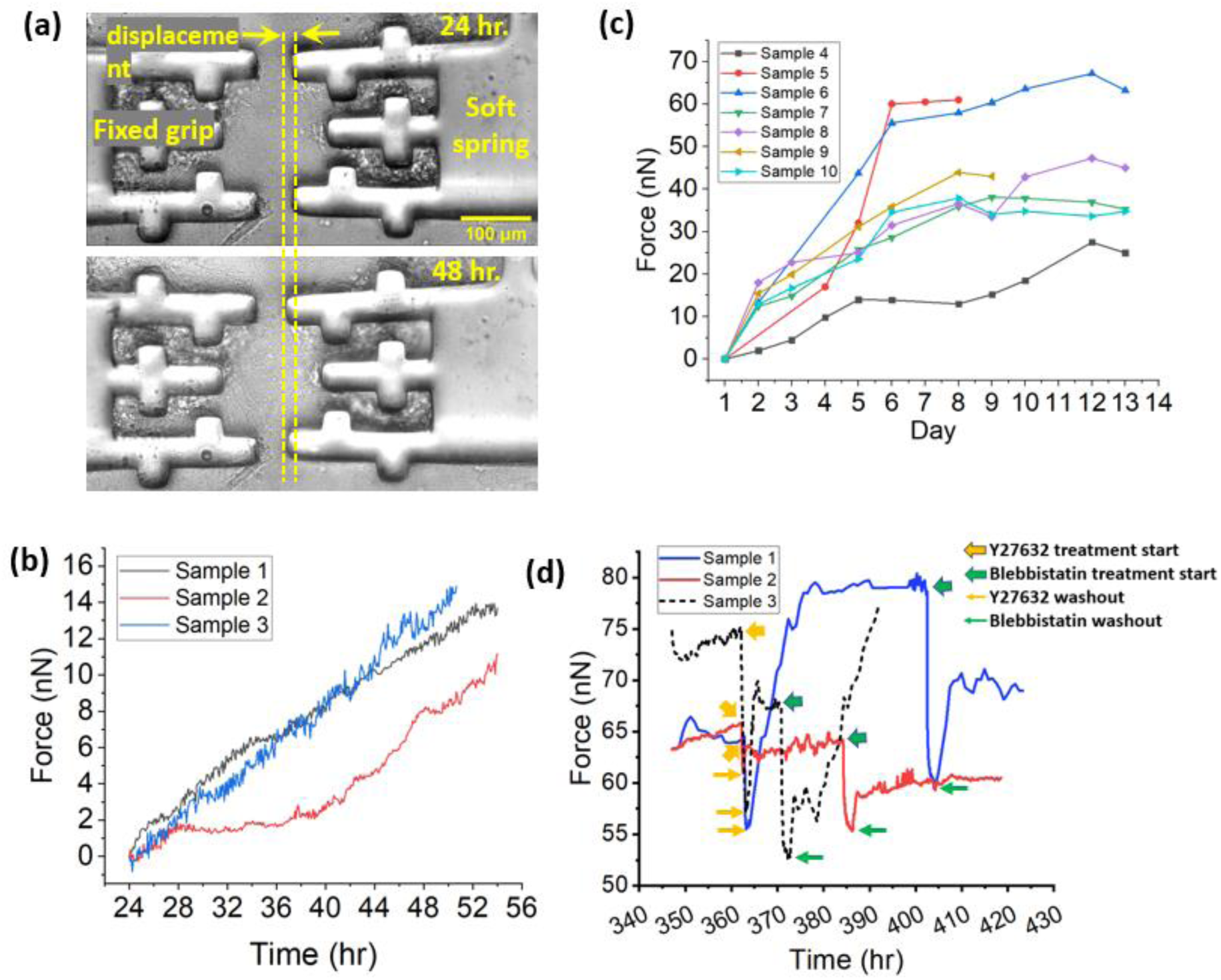
Neuronal network of rat hippocampal neuron continuously produce force and become saturated at around DIV13; (a) Bright field image of neuron on the force sensor showing the displacement of grip due to the force produced by the neuron; Time evolution of force produced by rat hippocampal neurons in 3D matrix (b) imaged every 5 min continuously (c) measured every day till DIV13; Myosin II inhibitor (y27632 and Blebbistatin) disrupts the force of neuronal network which is recovered upon washout (d) Measurement of force drop in 3D neuronal tissue due to Y27632 and Blebbistatin treatment

Our goal was to measure force solely generated by neurons and not by other types of cells-in particular glial cells, such as astrocytes. To achieve this goal, several measures were taken. Firstly, during tissue formation, only neuronal cell bodies were placed within the grips and the center area between the grips was kept free of cells. Our previous work ^23^ has already established that the force generated by cells inside the grip does not transfer to the force-sensing beam. Therefore, by keeping the center area free of cells (Materials and Method), we ensured that only neurites occupying the center region contributed to the force measured by the sensor, and any force generated by the cells inside the grips was not transferred to the force-sensing beam. Moreover, glial cell treatment was performed to minimize the presence of glial cells, following the published protocol by ^27^. This treatment significantly reduced the number of glial cells in rat hippocampal neuronal culture (Figure S1). As a result, the chance of having any force contribution from the glial cells would be minimized.

Secondly, the force generated by isolated astrocytes was compared to the neuronal force to explore any possibility of temporary migration of cells to the center area. The astrocyte force was significantly greater and had a much shorter time to force generation compared to the time constant of neuronal force (Figures S2, S3 and S4). The force produced by a single astrocyte was 25-30 nN (Figure S3), and in the case of multiple astrocytes, the maximum force was 200-300 nN (Figure S2). The time constant of astrocyte force for single and multiple astrocytes was ∼7.5 hrs. and ∼3 hrs. respectively, while the time constant for neuronal force was ∼3.5 days (∼83.4 hrs.) (Figure S4). This result indicated that a single astrocyte produced ∼20 nN force in only 3 hours, while the neuron force was ∼65 nN in 14 days. These imply that any contribution from astrocytes would appear as a large fluctuation in force measurements. Even if a single astrocyte migrated to the center, we would read a sharp rise in force within a few hours. No such fluctuations were observed in any of the neural network force measurements. Therefore, we concluded that there was no contribution of astrocyte force, and only neural network force was measured.

Thirdly, any possibility of force emerging from ECM polymerization (crosslinking) was tested by measuring the force produced by only ECM without the presence of any cell. The force was within the noise limit of the sensor (∼1 nN) (Figure S5), eliminating any force contribution from ECM in neuronal network force measurement. To further confirm that the force is produced by cells not ECM, we disrupted the force of neuronal tissue using Y27632 and Blebbistatin. Y27632 disrupts the ROCK pathway of force generation while Blebbistatin disrupts the ATPase activity of myosin II. We found that when neuronal tissue was treated with Y27632 and Blebbistatin, the force decreased by 7.4 - 23.5 % and 13.8 - 22.1% respectively (Figure 3(d) and Table 1). Upon washout of the drugs, the force was recovered within a few hours. The recovery of force after the washout of the drugs further supports the conclusion that the network force was produced exclusively by neurons, and not by the ECM. In summary, the force measured with our apparatus was due to neuronal contractions.

**Table1.**
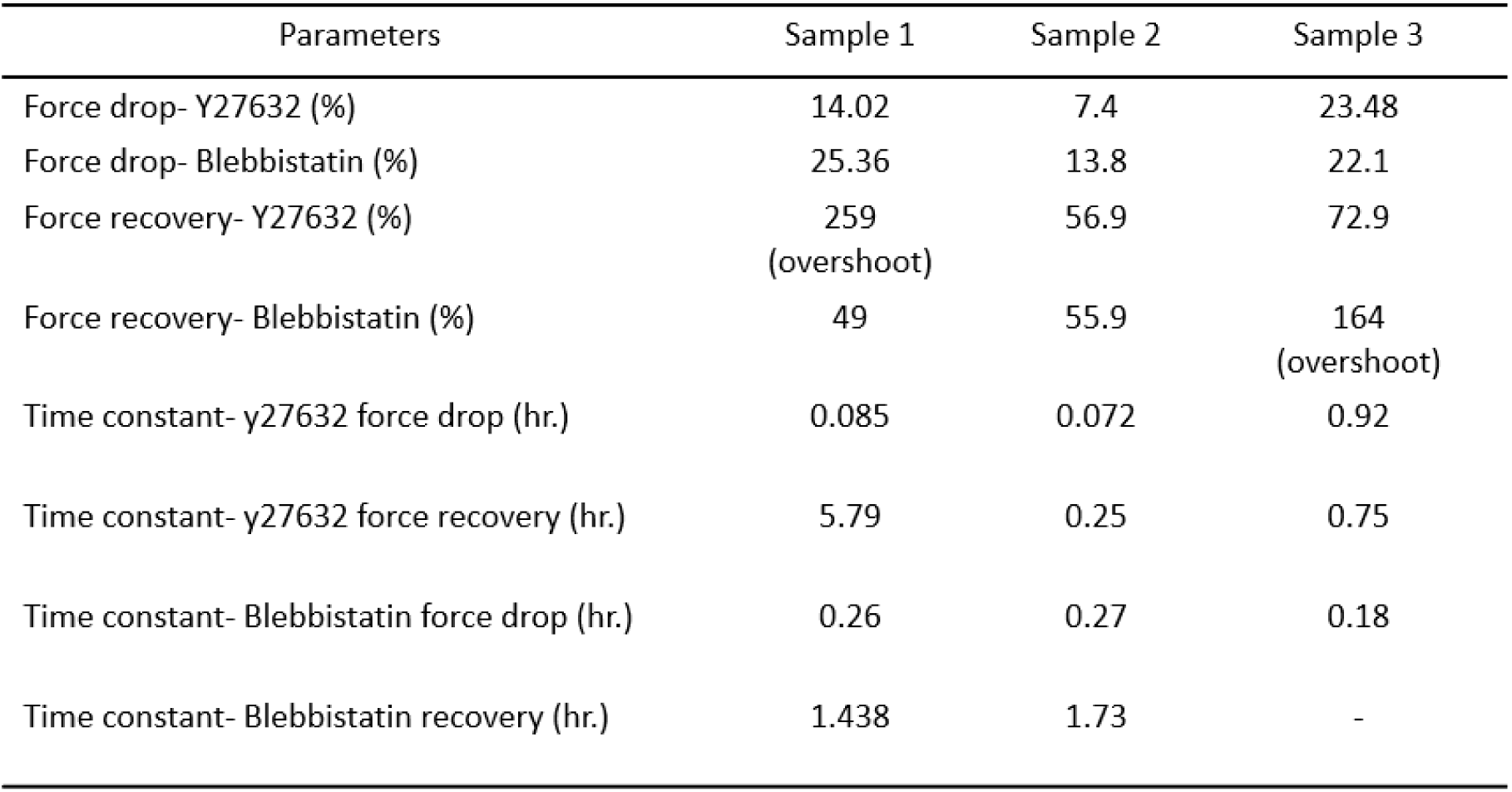
Force drop and recovery parameters for Y27632 and Blebbistatin treatment.

### Force relaxation disrupts neuronal firing

We now address the question of whether neuronal tension regulates the firing of the neuronal network.

We employed two different methods to measure neuronal firing and assess the impact of force disruption at the population scale (field potentials) and at the individual synapse scale. First, we used the Multi-Electrode Array (MEA, Multi-Channel Systems MCS GmbH) system to record the electrical activity of a population of neurons cultured on a 2D substrate with and without Blebbistatin; then we used the fluorescent protein, iGluSnFR, with and without Blebbistatin to measure the effect of force disruption on the firing of individual synapses. iGlusnFR is a genetically engineered fluorescent protein that has a sensitivity to glutamate.

To obtain the population-level response, hippocampal neurons were cultured on the MEA surface in neurobasal-based media (Gibco). At DIV 19, when the synapses become fully matured ^29^, we measured the firing rate of neurons before and after the treatment with Blebbistatin. The treatment with 1 hr. of Blebbistatin at 10 µM concentration, resulted in a decrease of over 60% in spike rate (Figure 4(a)), and a drop of approximately 90% in burst rate (Figure 4(b)). We then washed out the Blebbistatin solution, replaced it with neurobasal media, and measured neuronal firing after 24 and 48 hours. Interestingly, both spike and burst rates recovered after a 48-hour washout period. These findings suggest that mechanical tension is necessary for maintaining the firing of neurons at the population level, as neurons are unable to fire when the neuronal force is disrupted. The recovery of firing rates after the Blebbistatin washout further confirms the reversibility of the force disruption effect. In order to check whether the observed effect of force disruption extends to neurons from other tissue model systems, we conducted a similar experiment using primary mouse hippocampal neuron culture. A one-hour treatment with blebbistatin resulted in a significant decrease in spike rate and burst rate, which subsequently recovered after a 24-hour washout (Figure S7). These findings suggest that the impact of force disruption on neuronal firing is not limited to rat hippocampal neurons alone, other types of mammalian hippocampal neurons exhibit similar effects.

**Figure 4:**
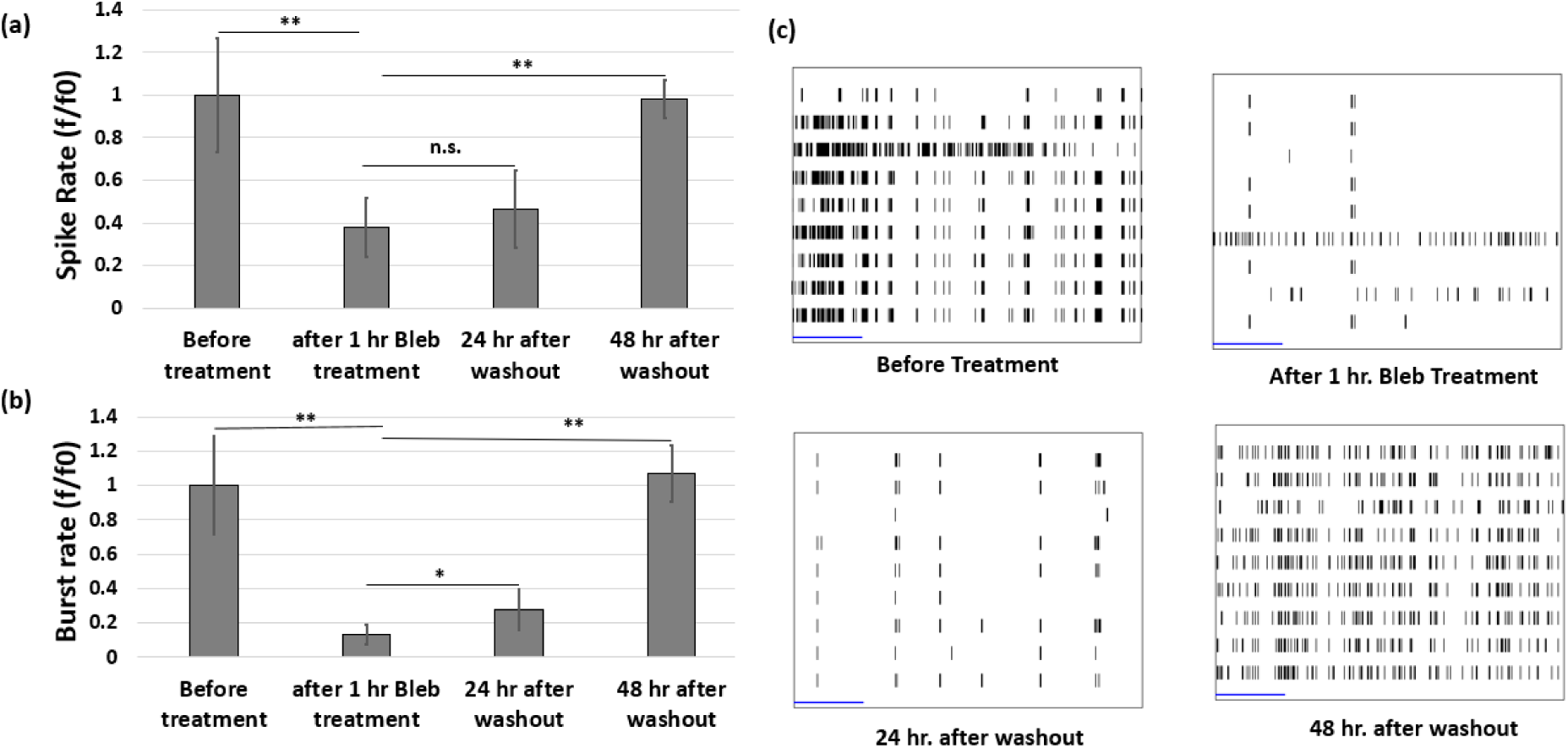
Force disruption using Blebbistatin significantly decreases the firing of the neuronal network (measured using MEA) which is recoverable after drug washout. Neuronal firing data from MEA. Change in (a) spike rate (n=6), and (b) burst rate (n=6) due to Blebbistatin treatment and the corresponding washout; (c) Raster plot before, during and after the Blebbistatin treatment. Statistical significance was determined using one-sided ANOVA with Tukey test; n.s., not significant; ∗p < 0.05; ∗∗p < 0.001; ∗∗∗p < 0.0001. Error bars indicate ± SEM.

While the MEA system provides information about the field potential of the neuronal culture recorded by the electrodes, it does not offer insights into individual synapses ^30^. To quantify the effect of force disruption on neuronal firing at the individual synapse level, we used iGluSnFR ^24^, a genetically engineered fluorescent protein. We transfected the 2D culture of rat hippocampal neurons with iGluSnFR which is expressed in the postsynaptic part, and it changes its fluorescence when it binds to the neurotransmitter glutamate. Since glutamate is the primary neurotransmitter released by hippocampal neurons during firing, a change in fluorescent intensity of iGluSnFR is an indicator of neuronal firing. We co-transfected the neurons with Lifeact or Homer, to identify the post-synaptic terminal ^31, 32^. An overlay image of iGluSnFR and Lifeact (Figure 5(a-c)) shows the dendritic spine along the length of the neurite. This helps to identify the position of individual synapses. We carried out time-lapse imaging (Figure 5(d)) (at 50 Hz) of the transfected neurites and captured their firing pattern. We identified the time of the iGluSnFR spike and calculated the rate. Figure 5(e) shows the firing pattern of individual synapses of this control group. For this group, we found that the average spike rate to be around 2.2 spike/sec (Figure 5(h)). To check the effect of force disruption on individual synapse firing, we used a 1-hour Blebbistatin treatment. Disruption of neuronal force significantly decreased (∼90% drop) the spike rate of the individual synapse (Figure 5(f) & 5(h)). The rate recovered after 24 hrs of washout of the drug (Figure 5(g) & 5(h)). Spike height distribution of neuronal firing from three groups, namely control, treatment, and treatment+washout (Figure S8) exhibits a similar pattern, characterized by several peaks in the height distribution. These peak positions are found to be multiples of the previous peak positions, indicating that glutamate release during neuronal firing follows a quantal process. The similarity of the pattern observed in all three groups suggests that the fundamental mechanism of glutamate release is not affected by blebbistatin treatment. However, it is possible that the number of vesicles available for release near the synaptic membrane has decreased due to the disruption of force caused by the treatment. These results indicate that indeed tension is necessary for neurons to function. To assess whether blebbistatin treatment has any detrimental effect on the health of neuronal culture, we performed a live/dead assay (Invitrogen). The results obtained from the assay (Figure S9) indicate that the viability of the neuronal culture remains unaffected by blebbistatin treatment.

**Figure 5:**
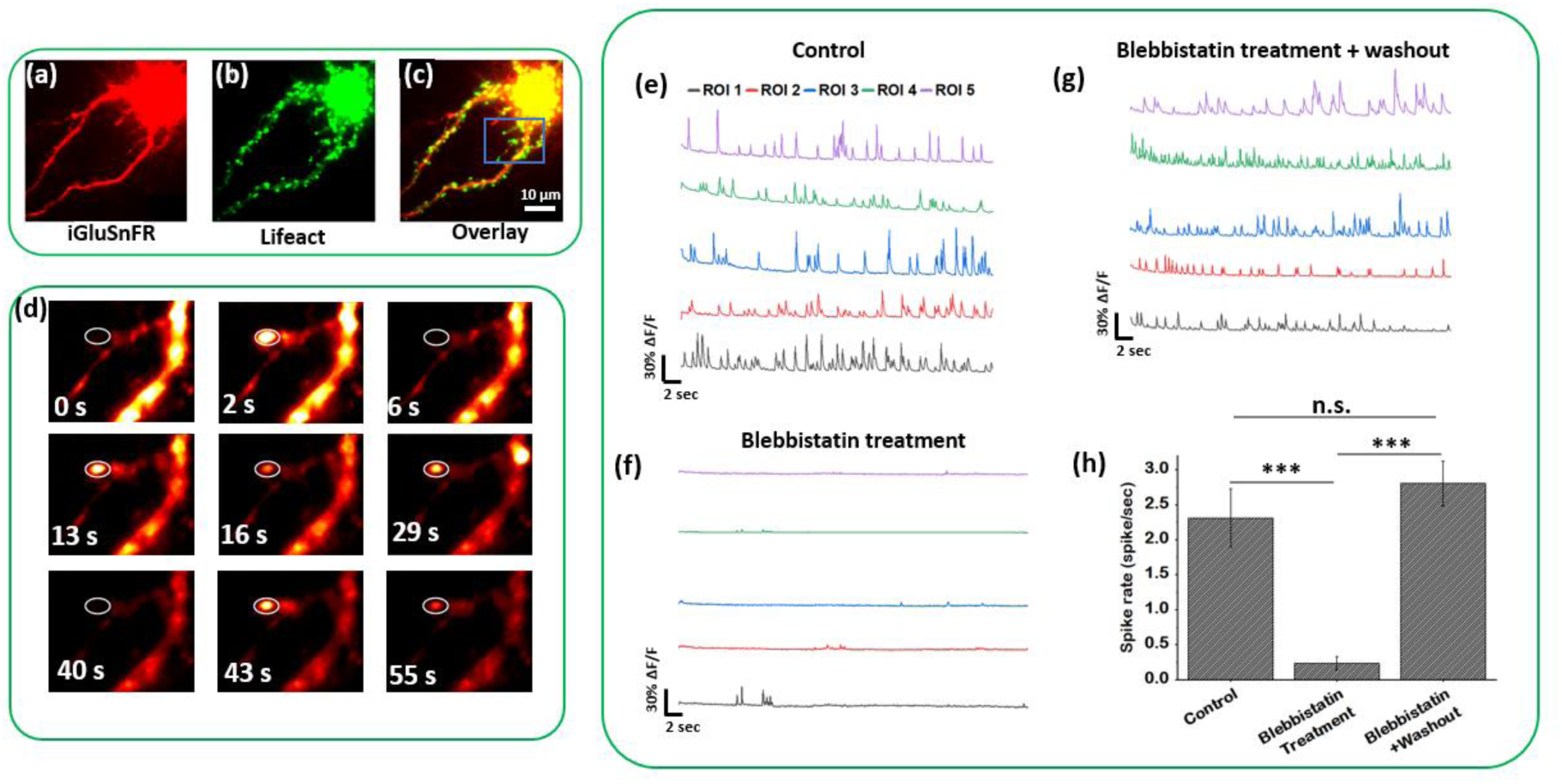
Force disruption using Blebbistatin significantly decreases the neuronal firing at single synapse level which is recoverable after drug washout; Rat hippocampal neuron co-transfected with (a) iGluSnFR and (b) Lifeact-miRFP; (c) Overlay image of iGluSnFR and Lifeact-miRFP showing the dendritic spine along the length of neurite; (d) Time series image of a single synapse (labeled by a white oval) showing the variation of iGluSnFR intensity at different time points; Time history of firing pattern at the individual synapse of neuronal culture from (e) control group; (f) Blebbistatin treatment; (g) Blebbistatin treatment with washout group. (h) spike rate at the individual synapse for the neuronal culture of different groups (n=3). Chart is showing mean ± SEM. Statistical significance was determined using one-sided ANOVA with Tukey test; n.s., not significant; ∗p < 0.05; ∗∗p < 0.001; ∗∗∗p < 0.0001.

### Force disruption decreases pre-synaptic vesicle intensity

Neuronal firing depends on the density of presynaptic vesicles (^33–36)^. Our results provide evidence that force disruption significantly decreases the firing rate. Therefore, we hypothesized that disruption of neuronal force may cause a decrease in presynaptic vesicles. To test this hypothesis, we applied Blebbistatin (10 µM) for 1 h on a 2D culture of rat hippocampal neurons on a petri dish at DIV14. Neurons were then fixed and stained for synaptophysin, a synaptic vesicle membrane protein ^37^. The intensity of synaptophysin, measured by confocal imaging, gives a measure of the amount of vesicle at the presynaptic terminal. Samples were co-stained with presynaptic marker Bassoon and post-synaptic marker Homer. Synapses were identified from the colocalization of these pre-and post-synaptic markers (Figure 6(a) & 6(b)). Only the positions where pre-and post-synaptic markers colocalize were considered as a synapse, and the intensity of the synaptophysin (vesicle) within these synapses was measured. We found that blebbistatin treatment causes a significant drop (∼20% decrease) in synaptic vesicle intensity (Figure 6(f)), suggesting that mechanical tension is crucial for maintaining vesicle clustering at the presynaptic terminal. However, after 24 hours of washout, vesicle intensity did not recover, indicating that it may take a longer time to replenish the lost vesicles.

**Figure 6:**
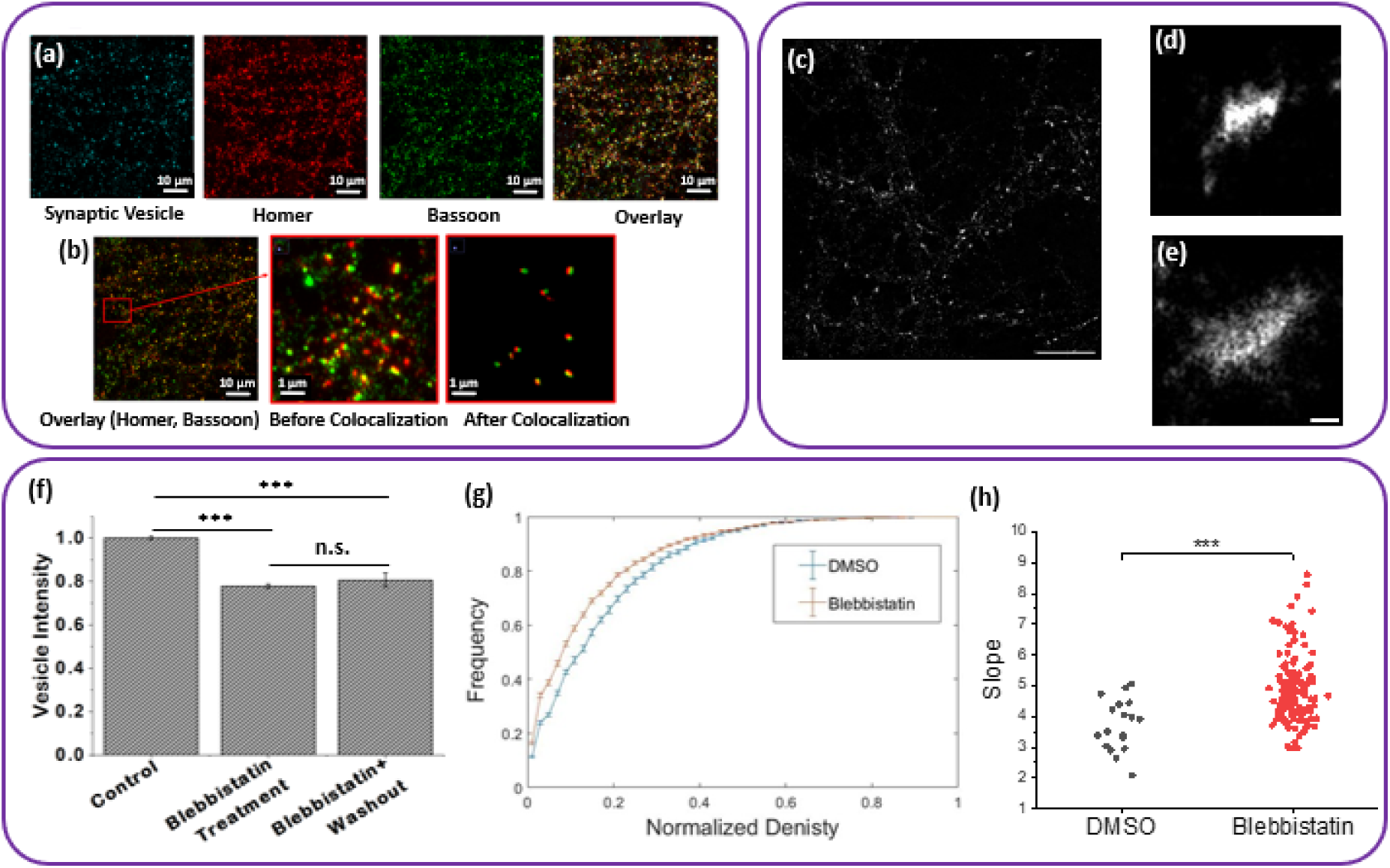
Force disruption using blebbistatin decreases pre-synaptic vesicle intensity. (a) Immunofluorescence image of the 2D culture of Rat hippocampal neuron. The sample is stained for Synaptic vesicle (synaptophysin) along with Post synaptic marker Homer and presynaptic marker Bassoon; (b) Overlay image of pre-synaptic and post-synaptic marker. Zoomed-in view shows the colocalization analysis technique used to identify the synapse. Only the positions where pre-synaptic and post-synaptic marker colocalizes are considered as a synapse; (c) STORM reconstruction of synaptophysin Alexa 647. (d,e) Inset of two selected synapses. Scale bar, 10um. Inset scale bar, 500nm. (f) Normalized intensity of synaptophysin at the synapse for the neuronal culture of different groups; (g) cumulative distribution plot of the pixel values of STORM reconstruction (normalized to maximum pixel value and averaged over synapses). DMSO (blue, n=19), Blebbistatin (orange, n=135). Error bars, standard error of the mean. (h) slope of the first four points of the synapse density cumulative distribution plots. two-way t-test; n.s., not significant; ∗p < 0.05; ∗∗p < 0.001; ∗∗∗p < 0.0001.

The loss of neuronal firing we observed under blebbistatin treatment motivated us to ask if this loss was due to a reduction in the density of synaptic vehicles. To answer this question, we performed STORM microscopy on fixed hippocampal neurons that were subjected to either DMSO or blebbistatin treatment (Figure 6(c-e)). To avoid counting artifacts due to variations in staining density from synapse to synapse, we chose to report the normalized density of each synapse. We plotted the normalized STORM reconstruction pixel values of each synapse as a cumulative distribution function (CDF) (Figure 6(g)). The CDF is a function of x that plots the integrated probability from 0 to the x-value (∫^*x*^ *p*(*x*)*dx*). At the maximum value of the distribution, the CDF has a value of 1 because all values have been integrated. As such, the slope of the CDF determines if a population is dominated by lower or higher values with a sharper slope meaning the distribution is dominated by lower values.

We noted a steeper rise in blebbistatin-treated CDF indicative of a higher frequency of pixel values with lower density. We quantified this by measuring the slope of CDF to find a significantly higher slope for blebbistatin-treated synapses (Figure 6(h)). This implies that blebbistatin-treated synapses have a smaller synaptic region populated by high synaptic vesicle density. This, together with confocal results, suggests that vesicle clustering reduced (20%) with force relaxation, and the remaining 80% of vesicles occupy a smaller space. It thus appears that blebbistatin causes 20% less vesicles in force-relaxed synapses. This is not to say the density is altogether lower for blebbistatin-treated synapses relative to control. A lower area of high synaptic vesicle density may be responsible for the loss of firing we observed.

## Discussion

In this study, we demonstrated that hippocampal neurons become contractile after forming synapses with each other. Our ultra-sensitive force sensor, with a resolution of 1 nN, allowed us to measure, for the first time, the evolution of force generation during the development of the neuronal network in 3D ECM.

We have also demonstrated, for the first time, that mechanical tension regulates the firing rate of rat hippocampal neurons. Using MEA (multi-electrode array) and iGlusnFR (a genetically encoded glutamate sensor), we showed that disruption of neuronal force with the drug Blebbistatin caused a 90% decrease in the spike rate of individual synapses. This result suggests a direct link between neuronal tension and the firing rate of a neuron. Our findings of mechanical tension regulating the firing rate of rat hippocampal neurons align with previous study ^38^ showing that motor protein myosin IIB plays a crucial role in maintaining dendritic spine morphology and excitatory synaptic transmission. Pharmacologic inhibition of myosin IIB with Blebbistatin resulted in a 40% depression of EPSCs amplitude. In comparison, our study found that disruption of neuronal force with Blebbistatin caused a 90% decrease in the spike rate of individual synapses. This larger decrease (90% vs 40%) in synaptic activity can be attributed to the differences in the experimental setup. Ryu et al. (2006) used rat hippocampal slices that freely floated during EPSC measurement, causing partial loss of tension before the Blebbistatin treatment. Conversely, we used a 2D culture of attached rat hippocampal neurons on a glass substrate, resulting in a higher tension before the Blebbistatin treatment.

Additionally, we studied the role of mechanical tension in presynaptic neurotransmitter vesicle clustering in rat hippocampal neurons. We found that disruption of force using Blebbistatin causes a ∼20% decrease in neurotransmitter vesicle clustering at the presynaptic terminal suggesting that mechanical tension is necessary for the clustering. This finding is consistent with an in vivo study showing that the axons of embryonic Drosophila motor neurons develop tension after forming neuro-muscular junctions (NMJ), and the tension is necessary for the clustering of neurotransmitter vesicles at the NMJ ^12^. Relaxing the tension by severing the axon using a laser beam (laser axotomy) resulted in the loss of vesicle clustering at the NMJ. However, restoring the tension by pulling the severed end of the axon using a micropipette, restored vesicle clustering. The implication is that mechanical tension might regulate vesicle accumulation at the synapse and that this regulation is conserved across different species.

It is puzzling to note in our experiments with hippocampal neurons that with blebbistatin treatment, the firing rate decreased by approximately 90% while the vesicle accumulation dropped by only around 20%. Further, after washing out of the blebbistatin, the neuronal firing rate was recovered in 24 hrs, but the vesicle clustering was not. One possible reason might be that blebbistatin treatment has two interdependent effects on the synapse. They both originate from a transition of vesicle dynamics within the synapse from processive to Brownian with the application of blebbistatin (^39^): (1) some of the vesicles from the reserve pool (RP) diffuse into the axon. These vesicles are conserved primarily through recycling by the kiss-and-run mechanism (^40^) after synaptogenesis is complete (^41^). If some of the vesicles are lost to the axon (about 20% in our experiments) by diffusion, then it may take a much longer time to be restored by the transport of vesicles from the cell body far away (100s of µm) from the synapse. This may explain the lack of recovery in vehicle clustering after the blebbistatin washout. (2) Blebbistatin disrupts synaptic vesicle transport (processive to Brownian ^39^) from the reserve pool (RP) to the close-by readily releasable pool (RRP) of the active zone site of the synapse. RRP consists of a small number of vehicles that are immediately available for release upon arrival of an action potential, whereas the RP is a larger pool of synaptic vesicles. The vesicles lost from the RRP are replenished by those supplied by RP through active transport ^42, 43^. In the presence of blebbistatin, firing decreases due to the depletion of vesicles from the active zone (RRP) and not being replenished from RP due to disruption of active transport. With washout of Blebbistatin, force is restored in a short span of time (30 mins, Figure 4), and the local transport from RP to RRP is restored, recovering the firing pattern within 24 to 48 hrs. A 20% decrease in RP vesicles, or 80% remaining reserve might have been sufficient to restore neuronal function in 24-48 hrs. Further investigation, using transmission electron microscopy, may help to support or refute this hypothesis, and further explain the differing effects of blebbistatin on the firing rate and vesicle clustering.

To summarize, we have successfully established a self-assembled 3D neuronal tissue platform where rat hippocampal neurons can grow and form synapses. This platform also allowed us to measure the evolution of force during the growth and development of the neuronal network with 1nN force resolution. Our results demonstrate that, much like the motor neurons in Drosophila fruit flies, rat hippocampal neurons also generate and maintain tension after synaptogenesis in vitro. This tension appears to be essential for neurons to carry out the basic function of firing and information transfer.

## Acknowledgment

Research reported in this publication was supported by the National Science Foundation (NSF) under award number 1935181. The content is solely the responsibility of the authors and does not necessarily represent the official views of the NSF.

## Author contribution

M.S.H.J. and M.T.A.S. conceived and designed the experiments. M.S.H.J., D.N.L., B.E., and K.Y.L. performed the experiments. M.S.H.J., D.N.L., B.E., K.Y.L., A.B., and M.A. performed imaging and analysis. S.R. performed vesicle intensity image analysis. M.S.H.J., D.N.L., P.S., and M.T.A.S. prepared the manuscript. All authors have read and approved the final manuscript.

## Competing interests

The authors declare that they have no competing interests.

## Data and materials availability

All data needed to evaluate the conclusions in the paper are present in the paper and/or the Supplementary Materials. Additional data related to this paper may be requested from the authors.

## Notes

### Competing Interest Statement

The authors have declared no competing interest.

