## Supplementary materials for "Synapses without tension fail to fire in an in vitro network of hippocampal neurons."

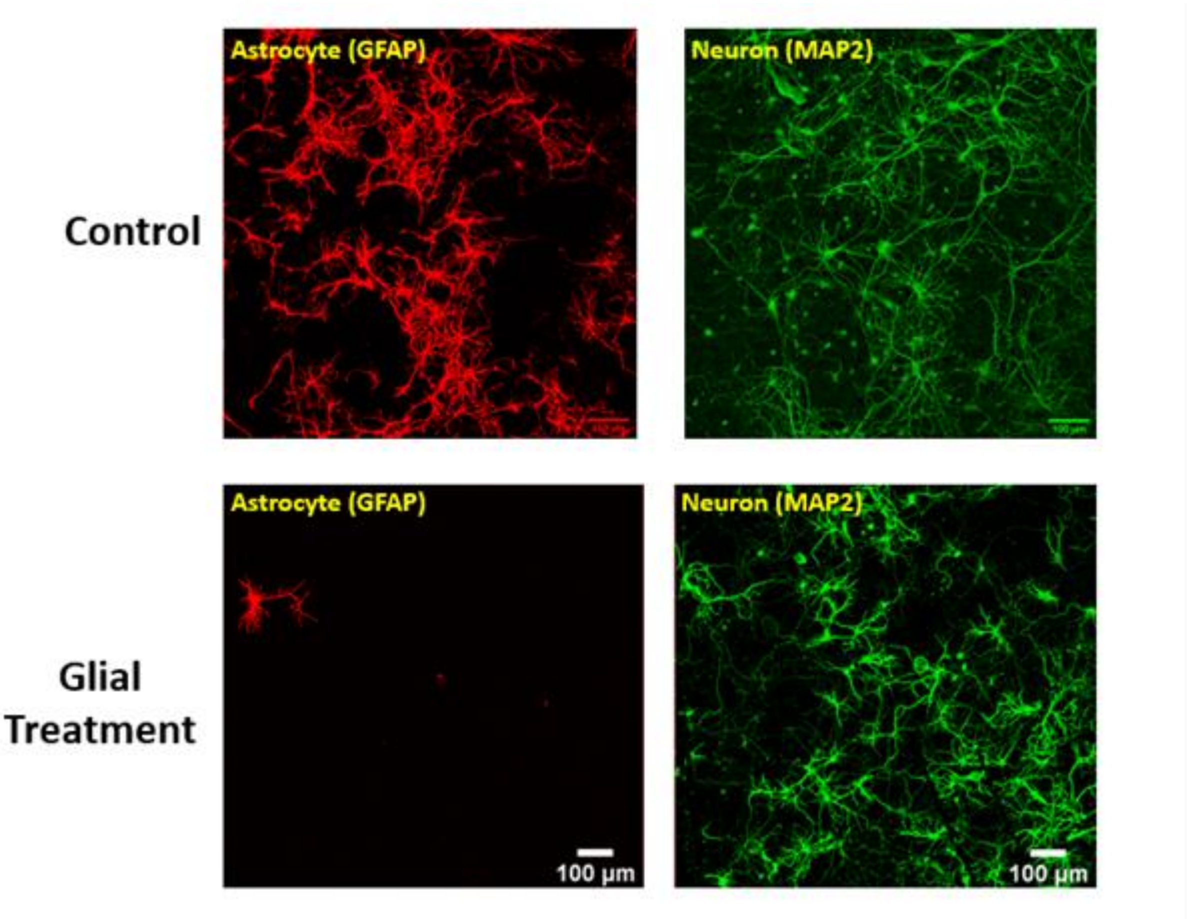


Figure S1: Effect of glial cell treatment on 2D culture of rat hippocampal neuron

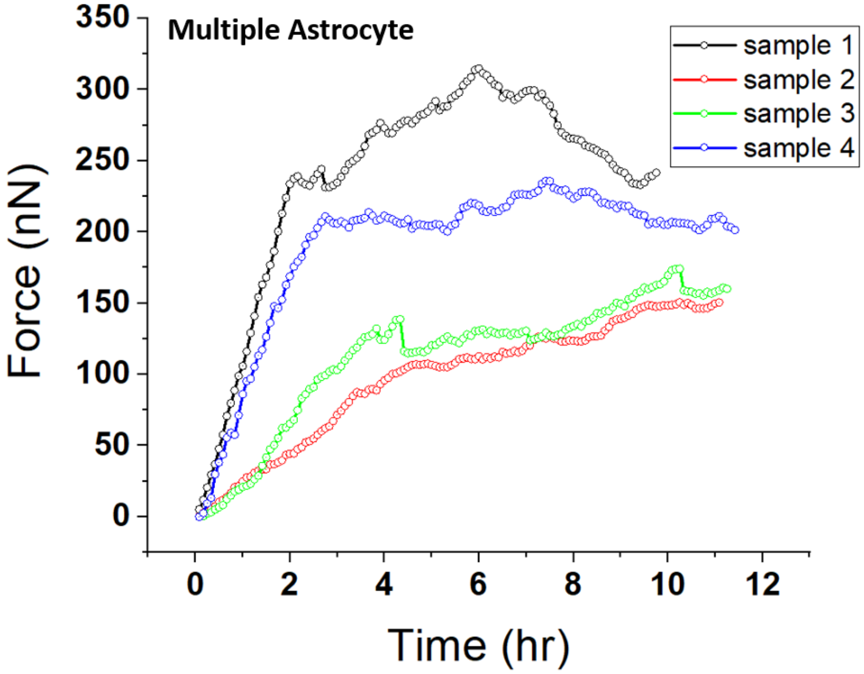


Figure S2: Time evolution of force produced by multiple astrocytes.

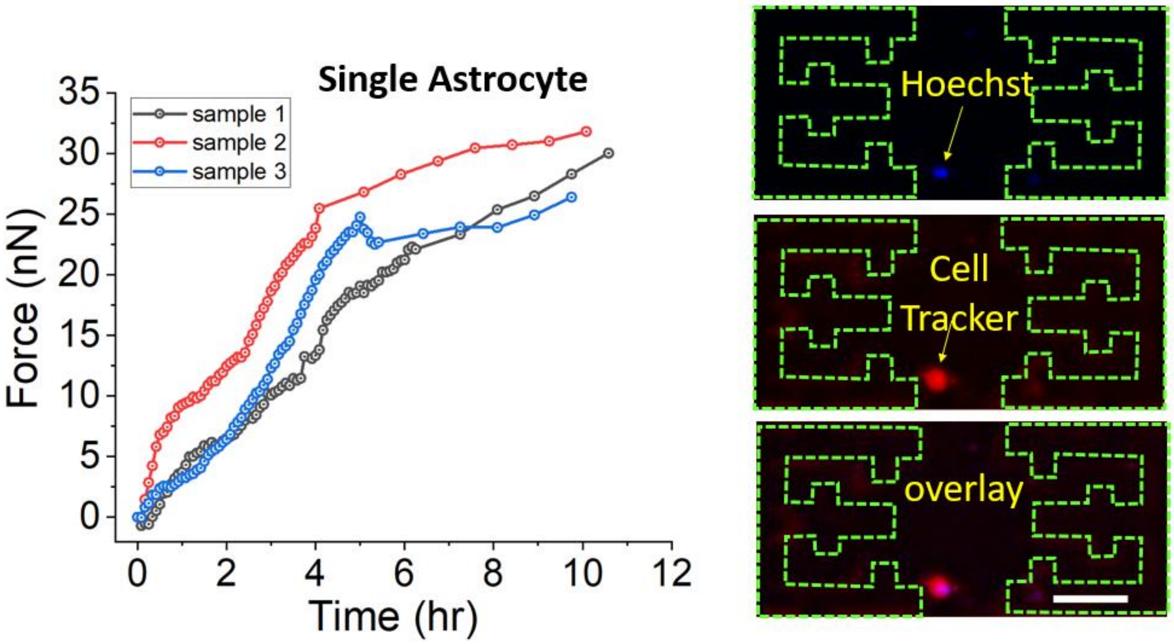


Figure S3: Time evolution of force produced by single astrocyte.

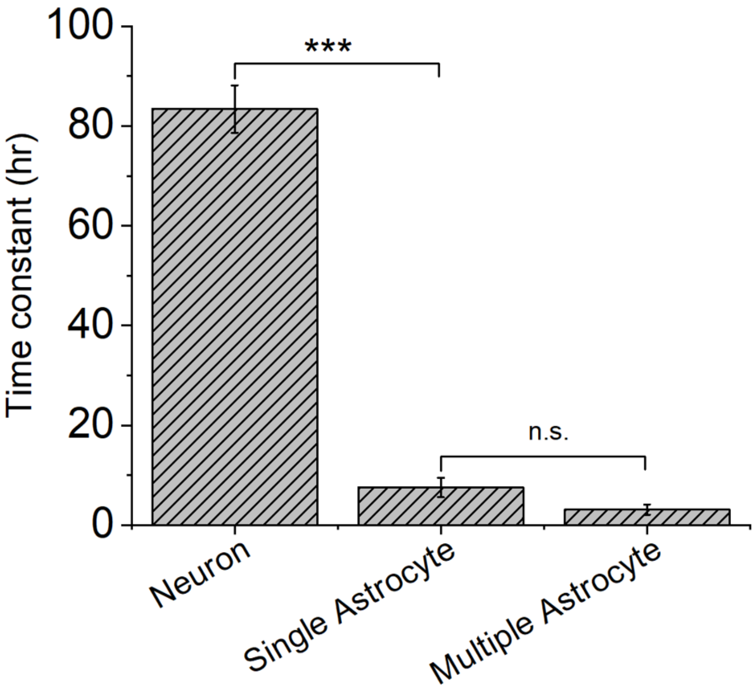


Figure S4: Time constant of force generation by single astrocyte, multiple astrocytes, and neurons

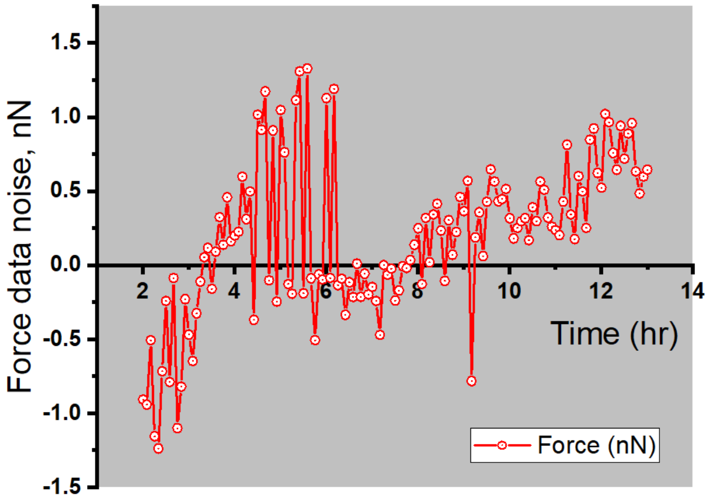


Figure S5: force data from sensor with tissue without any cell (noise data of force sensor)

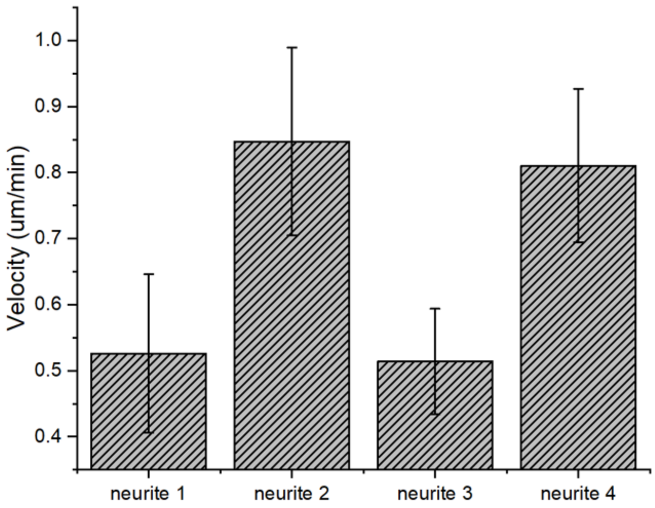


Figure S6: Instantaneous velocity of several growth cones cultured on the force sensor.

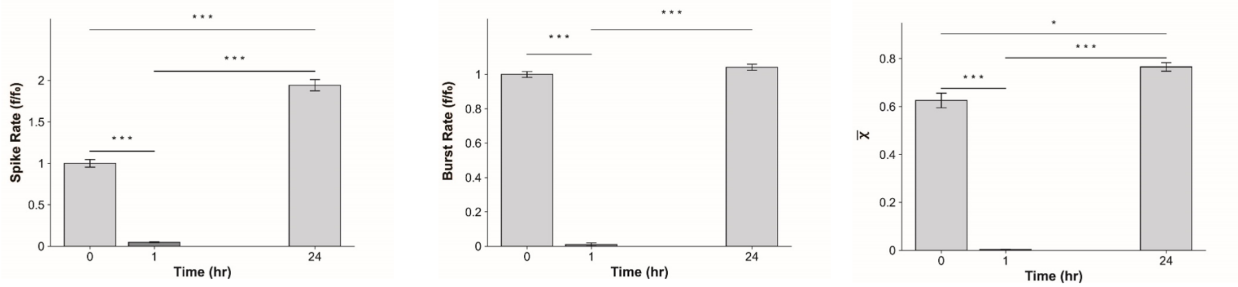


Time 0 hr → neuron in regular media at DIV11
Time 1 hr → recording after neuron was treated with 10 uM Blebbistatin for 1 hr
Time 24 hr → after Blebbistatin treatment neuron is washed out and recording is done after 24 hr

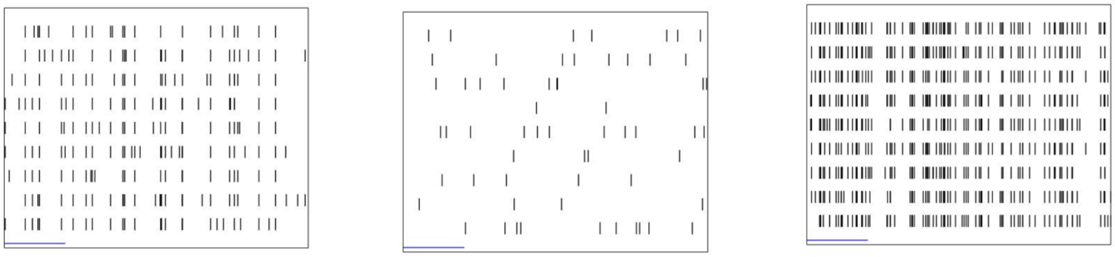

Before Blebbistatin. 2 s

During Blebbistatin

After washout

Figure S7: Force disruption using Blebbistatin significantly decreases the firing of mouse hippocampal neuronal network (measured using MEA) which is recoverable after drug washout.

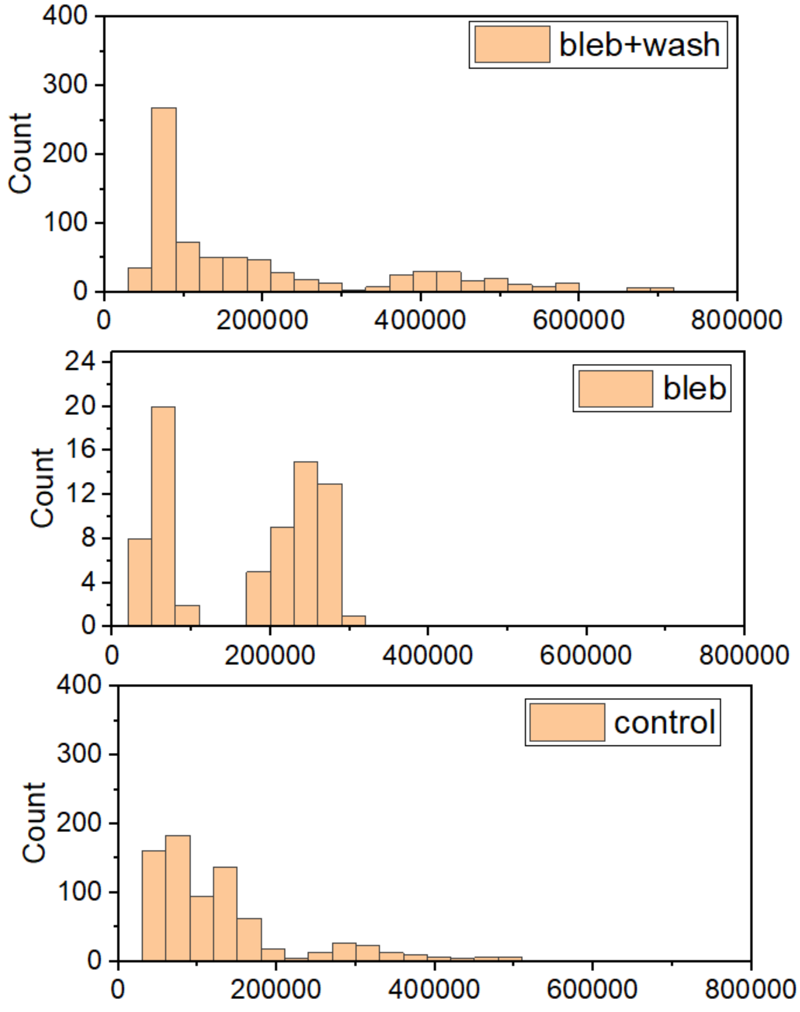


Figure S8: Histogram of neuronal firing spike height of neuronal culture from (a) control group; (b) Blebbistatin treatment; (c) Blebbistatin treatment with washout group.

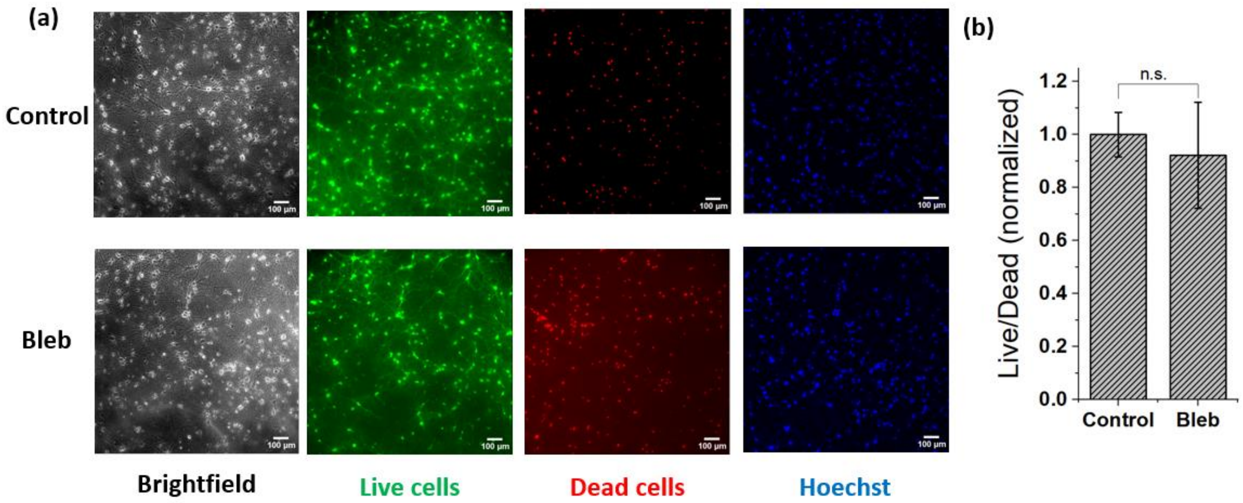


Figure S9: Result of live/dead assay showing that Blebbistatin treatment does not affect the viability of neurons.

supplementary video Legends -

Suppl. Video 1- Rat hippocampal neuron is growing and producing force on the force sensor platform.

Suppl. Video 2: Time lapse video showing the growth of rat hippocampal neuron. Filamentous actin (F-actin) of hippocampal neuron is labeled with SiR actin.

Suppl. Video 3: force produced by astrocyte in 3D matrix.

Suppl. Video 4: Force data with only ECM (no cell)

Suppl. Video 5: Neuronal firing measured with iGluSnFR.

Suppl. Video 6: Neuronal firing after 1 hr. Blebbistatin treatment
